# Automated Quantification of Enlarged Perivascular Spaces in Clinical Brain MRI across Sites

**DOI:** 10.1101/738955

**Authors:** Florian Dubost, Max Dünnwald, Denver Huff, Vincent Scheumann, Frank Schreiber, Meike Vernooij, Wiro Niessen, Martin Skalej, Stefanie Schreiber, Steffen Oeltze-Jafra, Marleen de Bruijne

**Affiliations:** Department of Radiology & Nuclear Medicine, Erasmus MC, the Netherlands; Department of Neurology, Otto-von-Guericke University Magdeburg, Germany; Faculty of Computer Science, Otto-von-Guericke University Magdeburg, Germany; Department of Neuroradiology, Otto-von-Guericke University Magdeburg, Germany; Department of Epidemiology, Erasmus MC, the Netherlands; Dept. of Imaging Physics, Faculty of Applied Science, TU Delft, the Netherlands; Center for Behavioral Brain Sciences (CBBS), Magdeburg, Germany; Department of Computer Science, University of Copenhagen, Denmark

**Keywords:** Perivascular Spaces, Deep learning, clinical MRI

## Abstract

Enlarged perivascular spaces (PVS) are structural brain changes visible in MRI, and are a marker of cerebral small vessel disease. Most studies use time-consuming and subjective visual scoring to assess these structures. Recently, automated methods to quantify enlarged perivascular spaces have been proposed. Most of these methods have been evaluated only in high resolution scans acquired in controlled research settings. We evaluate and compare two recently published automated methods for the quantification of enlarged perivascular spaces in 76 clinical scans acquired from 9 different scanners. Both methods are neural networks trained on high resolution research scans and are applied without fine-tuning the networks parameters. By adapting the preprocessing of clinical scans, regions of interest similar to those computed from research scans can be processed. The first method estimates only the number of PVS, while the second method estimates simultaneously also a high resolution attention map that can be used to detect and segment PVS. The Pearson correlations between visual and automated scores of enlarged perivascular spaces were higher with the second method. With this method, in the centrum semiovale, the correlation was similar to the inter-rater agreement, and also similar to the performance in high resolution research scans. Results were slightly lower than the inter-rater agreement for the hippocampi, and noticeably lower in the basal ganglia. By computing attention maps, we show that the neural networks focus on the enlarged perivascular spaces. Assessing the burden of said structures in the centrum semiovale with the automated scores reached a satisfying performance, could be implemented in the clinic and, e.g., help predict the bleeding risk related to cerebral amyloid angiopathy.

## 1 Introduction

Enlarged perivascular spaces (PVS) are structural brain changes visible on MRI. They can be identified as thin hyperintense tubular structures on T2-weighted MRI scans. PVS are increasingly thought to reflect the presence of cerebral small vessel disease, which represents a leading cause of cognitive decline and functional loss in elderly patients. In most studies, enlarged perivascular spaces are quantified using visual scores that either classify the burden of PVS in several categories [8], or count PVS [1]. These quantification methods are tedious and observer-dependent. Recently, several methods have been proposed to automatically quantify PVS burden [2, 4–6, 10, 13]. None of these methods have been evaluated in clinical scans, which present multiple challenges for the quantification of PVS. While in research studies, the scanning is highly standardized (same machine, same protocol, same scanning parameters, same investigators, etc.) to yield comparable results, this is not the case in clinical routine. The lower resolution of clinical scans also results in the computation of less accurate shape features, the most discriminative feature for the detection of PVS. Moreover, other MRI markers related to cerebral small vessel disease – such as white matter hyperintensities – are more prevalent in clinical scans than in population studies [5, 6, 4, 2] and could be confused with PVS because of their similar appearance.

In most studies, PVS are quantified separately in one or several clinically and epidemiologically relevant brain regions: midbrain, hippocampi, thalamus, basal ganglia, and centrum semiovale. In PVS research, the centrum semiovale is the most studied region, as PVS burden there has been most strongly associated to potential determinants of PVS and outcomes thereof. The centrum semiovale is also often the region with highest inter-observer agreement in the visual scoring of PVS [1]. In this study, we quantified PVS in the hippocampi, basal ganglia, and centrum semiovale.

Zhang et al. [13] automatically quantified PVS on 7T MRI scans. Boespflug et al. [2] proposed an automated quantification method combining image intensities and morphologic features from several MRI sequences. They evaluated their method in the centrum semiovale in research scans. Sudre et al. [10] proposed to use recurrent neural networks to detect PVS and lacunar infarcts in 16 subjects of a longitudinal study investigating the relationship between cardiovascular risk factors and brain health. Van Wijnen et al. [11] regressed intensity distance maps of PVS in the centrum semiovale using neural networks. Recently, Dubost et al. [4] proposed to quantify PVS burden in four brain regions – mid-brain, hippocampi, basal ganglia, and centrum semiovale – with neural network regressors trained with image level labels: the count of PVS in the target brain region. In research scans, the authors showed that they could reach a correlation between visual scores and automated scores similar to that of the inter-observer agreement in each region. They also found that associations between 20 determinants of PVS and visual PVS scores, and between the same determinants and automated PVS scores, were similar. The same authors [5] proposed to use a more advanced model (GP-Unet) for weakly supervised detection of enlarged perivascular spaces. This method estimates simultaneously the number of PVS and a high resolution attention map that can be used to detect and segment PVS. We decided to study the methods of Dubost et al. [4, 5] as the validation experiments with associations with clinical variables already brought them one step ahead of other methods for the application to clinical practice.

In this article, we applied and compared the two methods of Dubost et al. [4, 5] on 76 clincial MRI scans with a varying, low resolution acquired in the clinical routine of a hospital using nine different scanners and different protocols, while using models weights learned from high-resolution population study MRI scans acquired at another hospital in a highly controlled and standardized setting using a single scanner and protocol. The networks were not fine-tuned to the clinical data. For preprocessing, we used FSL packages instead of FreeSurfer parcellations of [4, 5] to segment the regions of interest. Finally, we show examples of attention maps of GP-Unet.

## 2 Datasets

### Training data

The training data consists of 1600 T2-weighted MRI scans from 1600 elderly participants in a population study: the Rotterdam Study [7]. Scans were acquired on a single 1.5T GE scanner, in a highly controlled and standardized setting. The scan resolution was 0.5×0.5×0.8 mm^3^. PVS were visually scored by a single rater in all scans in the hippocampi, basal ganglia and centrum semiovale, following the guidelines of Adams et al. [1].

### Evaluation data

The MRI data used for evaluation were gathered retrospectively from the Picture Archiving and Communication System (PACS) of University Hospital Magdeburg. MRI scans with visible signs of cerebral small vessel disease were selected. All selected patients had cerebral microangiopathy, and were diagnosed with at least one of the following: ischemic (i.e. lacunar) stroke or transient ischemic attack, spontaneous intracerebral hemorrhage, dementia (i.e. Alzheimers disease or vascular dementia), and epileptic seizures. Initially, 100 acquisitions from 100 different patients were collected. 24 Scans were excluded from the experiments either because FSL segmentation of the brain structures failed or because scans could not be rated visually, e.g. due to insufficient image quality caused by motion artifacts or presence of other pathologies such as extremely large lesions. This leaves a total of 76 scans for the study. Since the acquisitions have been obtained during the clinical routine, they present a considerable variance with respect to various image properties such as artifacts or image resolution. T1-weighted and T2-weighted MRI scans have been acquired with 9 different scanners. Two of these scanners, a 3T and a 1.5T from Philips, make up 66 of the 76 images. In total, there are three 3T-, four 1.5T- and two 1T-scanners. Three of them were Siemens (two 3T, one 1.5T), the rest were Philips machines. The time frame in which the data was acquired is almost 15 years and ranges from August 2004 until March 2019. The majority of the scans (43) has been acquired within the last 5 years of this period. The number of male and female patients was 46 and 30, respectively. Table 1 provides additional information about the data set. PVS were scored visually in the hippocampi, basal ganglia and centrum semiovale following the guidelines of Adams et al. [1]. Two raters scored PVS, the inter-rater agreement is reported in Table 2.

**Table 1.** Characteristics of the clinical dataset (minimum, maximum, mean and standard deviation)

|  | min | max | mean | std |
| --- | --- | --- | --- | --- |
| Patient age (years) | 35 | 89 | 71.39 | 9.32 |
| In-plane (axial) resolution ($\text{mm}^2$) | 0.39 | 0.68 | 0.45 | 0.04 |
| Resolution in z (mm) | 3.30 | 7 | 4.94 | 0.89 |
| Spacing between slices (mm) | 0.60 | 6.60 | 4.73 | 1.04 |

**Table 2.** Correlation between visual and automated PVS scores. Pearson correlations between the first rater and GP-Unet, CNN, and the second rater for each region. Correlations were all significant (p-value < 0.01). Significant correlations after Bonferroni correction are in bold.

|  | GP-Unet | CNN | R2 |
| --- | --- | --- | --- |
| Centrum Semiovale | **0.78** | **0.52** | **0.75** |
| Basal Ganglia | 0.31 | 0.25 | **0.56** |
| Hippocampi | **0.51** | 0.33 | **0.64** |

## 3 Methods

The target brain regions (hippocampi, basal ganglia, and centrum semiovale) are first segmented, masked and cropped. The result is then processed by trained convolutional neural networks that predict the count of PVS in each region. The neural networks were trained with high resolution MRI scans of a population study, but were used to predict PVS count in routine clinical scans of a hospital. The study was approved by the local ethics committee (No 28/16).

### 3.1 Preprocessing

To match the resolution of scans in the training set, all clinical scans were linearly interpolated to a resolution of 0.5×0.5×0.8 mm^3^.

Dubost et al. [4] used FreeSurfer parcellations to segment brain regions. FreeSurfer brain parcellation lasts usually several hours, which may prevent its use in clinical routine. In this study, we used instead FIRST and FAST algorithms from the FSL package [9] to segment brain regions from the T1 sequence in a matter of minutes. FIRST could compute segmentation of the basal ganglia and hippocampi. FAST was used to segment the white matter for the centrum semiovale region. Dubost et al. [4] also evaluated their method in the midbrain. As midbrain segmentation is not implemented in FSL, this region was excluded from the study. The T1 sequence was then rigidly registered to the T2 sequence using FSL FLIRT, and the segmentation labels were propagated from the T1 space to the T2 space.

Following the guidelines of Adams et al. [1] for visual scoring of PVS, Dubost et al. [4] quantified PVS in the centrum semiovale in the neighborhood of the slice located 1 cm above the top of the lateral ventricles. As FSL does not compute ventricle segmentation, we used instead the segmentation of the basal ganglia as approximation, and selected the slice 1 cm above the top of the caudate nucleus.

The following preprocessing steps were computed exactly as described by Dubost et al. [4]. Namely, the segmentation masks were dilated, convolved with a gaussian kernel to smooth the border of the mask, and multiplied pixelwise with the T2 intensities. The masked regions were then cropped, normalized between 0 and 1 using the minimum and maximum intensity values in the masked region, and given as input to the neural networks.

### 3.2 Neural Networks

The preprocessed images were given as input to two different types of neural networks proposed for automated PVS quantification: (1) a neural network with four convolutional layers and a max-pooling layer which outputs the number of PVS in a region [4] and that we call *CNN*, and (2) *GP-Unet*, a similar neural network proposed by the same authors [5], in which the downsampling path is followed by an upsampling path to enable weakly supervised detection of PVS. Networks of both methods were trained with only image-level labels.

Attention maps of GP-Unet were computed to visualize the focus of the networks using a linear combination of the feature maps of the last convolutional layer, as described by Dubost et al. [5].

## 4 Results and Discussion

Table 2 shows the Pearson correlation, and Table 3 the mean absolute error, between visual and automated PVS scores for each region and for each method, and the corresponding inter-rater agreement. Scatter-plots are shown in Figure 1. Attention maps of GP-Unet are displayed for each region in Figure 2.

**Table 3.** Mean absolute errors between visual and automated PVS scores. Mean absolute error between the first rater and GP-Unet, CNN, and the second rater for each region.

|  | GP-Unet | CNN | R2 |
| --- | --- | --- | --- |
| Centrum Semiovale | 5.58 | 6.39 | 4.67 |
| Basal Ganglia | 5.67 | 5.49 | 3.78 |
| Hippocampi | 2.58 | 3.0 | 2.08 |

**Fig. 1.**
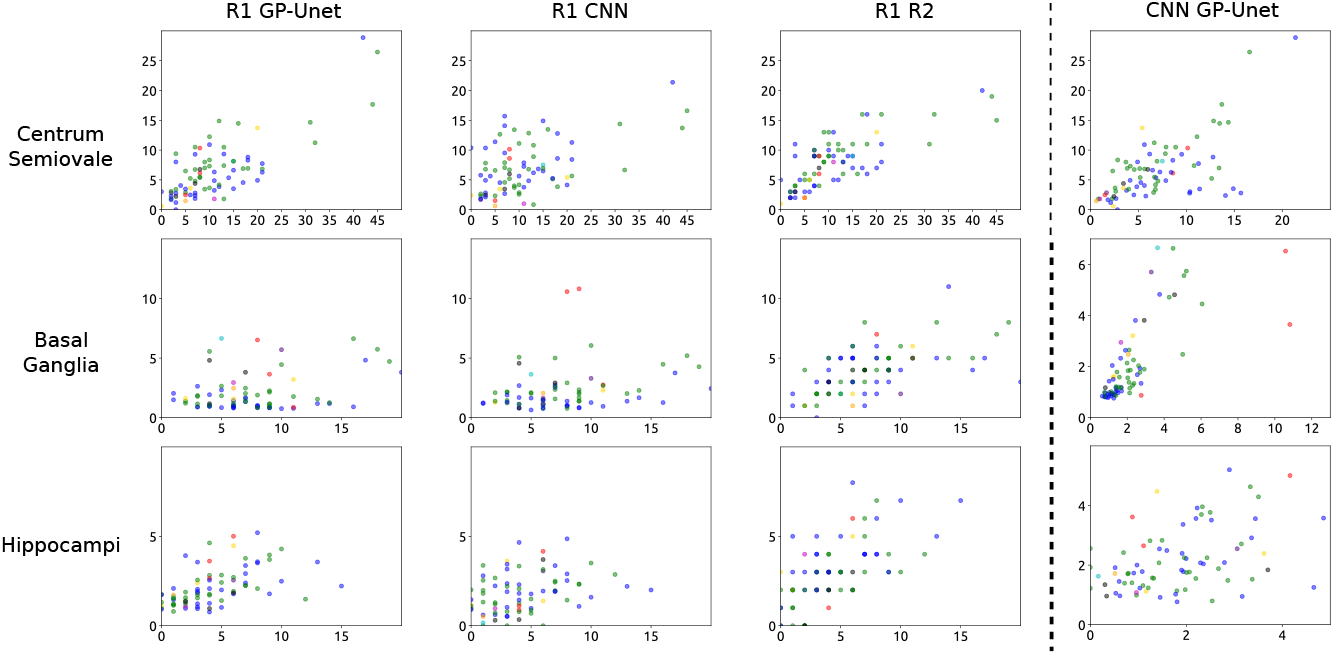
Comparison between visual and automated PVS scores. The different colors represent different scanners. The visual PVS scores of the first rater (R1), on the x-axis, are compared with the predictions of GP-Unet, CNN, and with the visual scores of the second rater (R2), on the y-axis. In the right column we plotted the automated PVS scores of GP-Unet versus those of CNN.

**Fig. 2.**
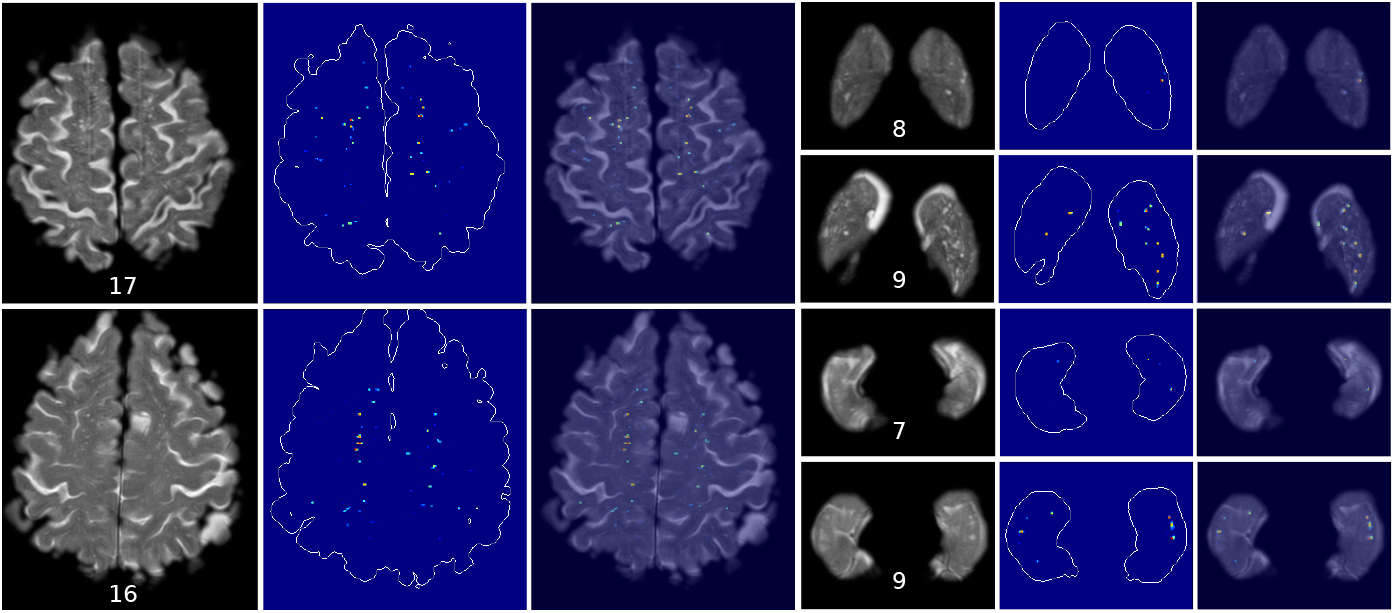
Attention Maps of GP-Unet in an axial view. Attention maps for the centrum semiovale are displayed on the left, for the basal ganglia on the top right, and for hippocampi, on the bottom right. Visual scores are indicated below each region. For each selected image, from left to right, we show the original image, the attention map with drawn contours of the region, and the overlay of both. The colormaps of the attention maps were manually adjusted for each image. Highlighted structures are considered as PVS by the networks. The redder a structure is, the higher is its weight in the computation of the automated PVS scores by the network. For the centrum semiovale, we selected two images that correspond to an average agreement between automated and visual score (human rater R1). For the basal ganglia and hippocampi, we selected one image with poor agreement (top), and another image with good agreement (bottom).

There was no noticeable difference in the computation of the regions of interest when using FSL masks instead FreeSurfer masks, but the interpolation to 0.5×0.5×0.8 mm^3^ was needed to reuse the networks optimized on high resolution scans. The visual PVS scores were highly correlated to the automated PVS scores of GP-Unet in the centrum semiovale (0.78 Pearson correlation), were moderately correlated in the hippocampi (0.52), and a lower correlation in the basal ganglia (0.28). Attention maps of GP-Unet (Figure 2) show that, as expected, the method focuses on perivascular spaces.

While on research scans, CNN and GP-Unet reached a similar performance in all regions, our experiments on clinical scans show that the correlation between visual PVS scores and automated PVS scores of GP-Unet was significantly higher than that of visual PVS scores and automated scores of CNN in the centrum semiovale (Williams’ test, p-value < 0.0001) and in the hippocampi (p-value < 0.05). Contrary to CNN, GP-Unet combines features of different scales via skip connections, which may have assisted the computation of discriminative shape features, and improved the detection of single PVS, as opposed to detecting – or missing because of their too large size – a cluster of PVS without being able to individually count them.

The correlation in the basal ganglia (0.31 for GP-Unet) is lower than in the other regions and is notably lower than the inter-rater agreement (0.56). Attention maps (Figure 2) show that the network only detects the largest PVS in the basal ganglia, and misses less enlarged PVS. The scatter-plots (Figure 1) seem to confirm this observation: in the basal ganglia, the networks underestimate the number of PVS, and predict similarly low numbers of PVS for all scans.

Table 2 shows lower inter-rater agreement for the basal ganglia than for the other regions. This might be a consequence of PVS being visually rated only in a single slice in this region [1]. The low resolution of clinical scans in z direction might cause a large variability in the selection of this slice, which might negatively influence the reproducibility of the visual rating. The automated methods quantify PVS in the complete volume of the basal ganglia, which was previously shown to be more reproducible than the visual PVS scores [6]. Interestingly, the automated PVS scores of both methods – CNN and GP-Unet – are highly correlated in the basal ganglia (0.73 Pearson correlation). The correlation between their scores was higher in the basal ganglia than in other regions.

Results in the centrum semiovale (0.78 Pearson correlation) are similar to the inter-rater agreement (0.75). This is also close to the inter-rater agreement (0.80 intraclass correlation coefficient) as reported in earlier studies in high resolution research scans [1]. Demonstrated quantification of PVS burden in the centrum semiovale could aid in the better stratification of cerebral small vessel disease subtypes, i.e. hypertensive arteriopathy and cerebral amyloid angiopathy, especially in large and hospital-based cohorts. This would presumably have important therapeutic and prognostic implications in terms of prescribing oral anticoagulants and preventing intracerebral hemorrhage. This is of particular importance in cerebral amyloid angiopathy, that has not only been related to severe PVS burden in the centrum semiovale [3], but also to a significantly higher risk for intracerebral bleeding in face of oral anticoagulant treatment. [12].

In future work, the results in the basal ganglia and the hippocampi may be improved by fine-tuning the neural networks using the clinical dataset, and by adding data augmentation during training with research scans to imitate the resolution of clinical scans and contrast variations between different scan protocols or scanners. The results presented are already promising considering the large differences between training and test sets.

The complete computation of the automated PVS scores lasts only a few minutes on CPU. Most of the computation time is spent on FSL brain structures segmentation and registration from the T1-weighted scans to the T2-weighted scans. After this preprocessing, the computation of the automated PVS scores took only about 6 seconds per brain region on CPU. This low computation time can facilitate the implementation of such a method in clinical practice.

## 5 Conclusion

We showed that PVS burden could be automatically quantified in the centrum semiovale in clinical scans, with an agreement with visual scores that was similar to the inter-observer agreement. Automated PVS scores were computed with a neural network that was trained high-quality research scans and with only global labels of PVS burden. These results could contribute to bringing automated PVS quantification to the clinic and guide the administration of anti-coagulant drugs.

## Acknowledgements

This work received funding from the Netherlands Organisation for Health Research and Development (ZonMw - Project 104003005) and the federal state of Saxony-Anhalt, Germany (Project I 88).

